# Classification of LTR Retrotransposons via Interaction Prediction

**DOI:** 10.1101/2024.02.11.579858

**Authors:** Silvana C. S. Cardoso, Douglas S. Domingues, Alexandre R. Paschoal, Carlos N. Fischer, Ricardo Cerri

## Abstract

Transposable Elements (TEs) are genetic sequences that can relocate within the genome, thus promoting genetic diversity. Classifying TEs in eukaryotes involves a hierarchy formed by classes, subclasses, orders, superfamilies, families, and subfamilies. According to this taxonomy, LTR retrotransposons (LTR-RT) constitute an order. The primary objective of this study is to explore the classification of LTR retrotransposons at the superfamily level. This was achieved by predicting interactions between LTR-RT sequences and conserved protein domains using Predictive Bi-Clustering Trees (PBCTs). Two datasets were used to investigate the relationships among different superfamilies. The first one comprised LTR retrotransposon sequences assigned to Copia, Gypsy, and Bel-Pao superfamilies, whereas the second dataset included consensus sequences of the conserved domains for each superfamily. Therefore, the PBCT decision tree tests could relate to both sequence and class attributes. In the classification process, interaction is interpreted as either the presence or absence of a domain in a given LTR-RT sequence. Subsequently, this sequence is classified into the superfamily with the highest number of predicted domains. Precision-recall curves were adopted as evaluation metrics for the method, and its performance was compared to some of the most commonly used models in the task of transposable element classification. Experiments on *D. melanogaster* and *A. thaliana* showed that PBCTs are promising and comparable to other methods, especially in the classification of the Gypsy superfamily.

## 1. Introduction

The unprecedented increase in the volume of biological data makes the development of effective analysis methods crucial. Transposable Elements (TEs) are one example of biological components that offer valuable insights into organisms from both practical and evolutionary perspectives (Schietgat *et al*. 2018).

TEs are genetic sequences that have the ability to move from one location to another within the genome, thereby promoting genetic variability. This phenomenon can have both evolutionary benefits and detrimental effects on the organism (Goerner-Potvin & Bourque 2018). TEs constitute significant portions of the genetic material in various species, particularly eukaryotes. This is the case with the human genome, of which approximately half corresponds to TEs (Goerner-Potvin & Bourque 2018).

The TE classification system proposed by Wicker *et al*. (2007) for eukaryotic organisms includes the levels of class, subclass, order, superfamily, family, and subfamily. Since classification is a task of great relevance in the study of transposable elements, strategies that utilize machine learning are promising. The class level is composed of: Class I or retrotransposons and Class II or DNA transposons. Class I corresponds to TEs that use reverse transcription as the basis for their transposition; a process in which the sequence is transcribed from DNA to RNA and then from RNA to DNA. For this reason, this process is usually associated with the “copy and paste” strategy. Class II does not depend on reverse transcription, and its transposition is based on a “cut and paste” strategy.

Class I can be divided into five orders: LTR retrotransposons (long terminal repeats (LTR) retrotransposons), DIRS-like elements (Dictyostelium intermediate repeat sequence - like elements), PLEs (Penelope-like elements), LINEs (long interspersed nuclear elements) and SINEs (short interspersed nuclear elements). Class II is divided into two subclasses distinguished by the number of DNA strands that are cleaved during transposition. Subclass 1 consists of TEs of the TIR order, which are characterized by their terminal inverted repeats (TIRs) of varying lengths. Subclass 2 groups the Helitron and MaverickOs orders.

The superfamilies of an order are defined through uniform characteristics that are widely spread, such as the protein structure or non-coding domain (Wicker *et al*. 2007). LTR retrotransposons are characterized by long repetitive termini, with the coding region flanked by DNA sequences that are typically homologous and range from a few hundred to 5kb (Schietgat *et al*. 2018). These elements are ubiquitous in plants, where they can comprise up to 95% of the genome, and also occur in substantial numbers in animals, where they remain active in species such as fruit flies and mice. Although inactive in humans, LTR retrotransposons constitute approximately 10% of their genome (Goerner-Potvin & Bourque 2018). According to Wicker *et al*. (2007), LTR retrotransposons are classified into five superfamilies with similar transposition mechanisms, namely Copia, Gypsy, Bel-Pao, Retrovirus and ERV. The Copia and Gypsy superfamilies are the most abundant outside the animal kingdom, and the Bel-Pao superfamily is structurally similar to them (Wicker *et al*. 2007).

According to Orozco-Arias *et al*. (2019), the classification of TEs is a complex task due to various factors, such as their dynamic evolution, which includes, for example, insertions of other TEs into a pre-existing TE and non-homologous recombination. For this reason, different methods have been developed that focus on the classification of a specific group of transposable elements. However, such methods have limitations, such as considering TEs with a predefined structure, which makes them insensitive to elements outside that standard structure.

In (Schietgat *et al*. 2018), a machine learning-based framework was developed for the identification and classification of transposable elements at the superfamily level within a given order. The application of this framework to LTR retrotransposons showed promising results. However, the classifier used was a conventional random forest, and the attributes of the problem classes and their correlations were not considered. This can be addressed by treating the classification task as an interaction prediction problem. In the literature, there are initiatives in this direction, although none of them focused on TEs. As an example, in (Santos *et al*. 2019), the concept of Predictive Clustering Trees (Vens *et al*. 2008) is extended and the Predictive Bi-Clustering Tree (PBCT) method is proposed, whose main feature is to work with attributes of both examples and classes.

In this context, the main objective of this work is to study the classification problem of LTR retrotransposons at the superfamily level as an interaction prediction task applying the predictive bi-clustering tree method. To this end, the classification strategy used was the detection of conserved domains, representatives of each superfamily, in the genetic sequences of TEs. As a way of exploring the correlations between the different classes considered (conserved domains) two datasets were used to train the model. The first was composed of sequences from TEs of the order LTR retrotransposons and the second by consensus sequences of the conserved domains of each superfamily. Therefore, the tests implemented in the decision tree could refer to both the set of attributes of the sequences and the set of classes (superfamilies).

The remainder of this paper is organized as follows. In Section 2 a bibliographical review of works proposed for the classification of transposable elements, more specifically LTR retrotransposons at the superfamily level, is performed. Section 3 presents the different approaches used in the literature for predicting interactions. Section 4 details the PBCT method used in our experiments, while Section 6 presents our proposal to use PBCTs to classify LTR retrotransposons. The experiments carried out and the analysis of their results are presented in Section 7. Finally, conclusions and future works are presented in Section 8.

## 2. Related Work

The LTRDigest method (Steinbiss *et al*. 2009) was designed specifically for Long Terminal Repeats (LTR) retrotransposons. It annotates LTRs with protein domains using Hidden Markov Models (HMMs) and also incorporates other structural features. LTRDigest can be used for classification by grouping LTR retrotransposons without any pre-defined classification scheme. To evaluate whether the created groups correspond to LTR superfamilies, Steinbiss *et al*. (2009) compared representative sequences from each group to a reference set composed of known transposable sequences.

TEClass is a method proposed by Abrusán *et al*. (2009) that uses a hierarchy of binary Support Vector Machines and k-mers as attribute vectors. The method classifies transposable elements into Classes I and II. The elements belonging to Class I can be further classified into LTRs and non-LTRs. The non-LTRs are then classified into SINE or LINE orders.

Feschotte *et al*. (2009) proposed a method called RepClass for classifying transposable elements. RepClass consists of three separate classification modules, each based on a different approach: homology, detection of structural features, and identification of target-site duplications. Each module produces a classification result at a different level, usually at the subclass or order level. Finally, the results from all three modules are combined to generate a single overall classification ranking.

The RepeatMasker method (Smit *et al*. 1996-2010) has a wide applicability for finding and masking repeats in sequences based on their similarity with annotated library sequences. It is widely used in the identification of TEs. The Censor method (Jurka *et al*. 1996), which is similar to RepeatMasker, uses the BLAST tool (Altschul *et al*. 1990) for comparison. Additionally, it removes overlaps and fragmented repeats.

The Pastec method (Hoede *et al*. 2014) uses different features for TE classification, including structural features (sequence length, presence of LTRs), homology, and conserved functional domains obtained from HMMs. Pastec generates classifications at the order level and includes all orders defined by the hierarchical classification established by Wicker *et al*. (2007).

In (Loureiro *et al*. 2013), machine learning techniques were used to improve TEs identification. Traditional methods were independently used to perform identification, and a classifier was built to combine these predictions and determine whether a sequence is a TE or not. Another classifier was applied to predict the best method for determining the exact boundaries of a TE.

In (Santos *et al*. 2018), several machine learning classifiers are used to explore different strategies for selecting positive and negative training instances and to apply local hierarchical classification methods for TEs classification.

The TE-Learner (Schietgat *et al*. 2018) is a versatile framework that uses Machine Learning to identify TEs in a given genome and classify them at the superfamily level of a given order. When applied to LTR retrotransposons, TE-Learner showed better predictive accuracy than the methods RepeatMasker, Censor, and LTRDigest. Furthermore, it was able to identify TEs that none of these three methods could detect.

Several works focusing on Deep Neural Networks (DNNs) for TE classification have been published, showing promising results. In Nakano *et al*. (2018), fully connected neural networks (FCNNs) were used to improve the hierarchical classification performance of TEs. FCNNs are a type of artificial neural network in which the architecture ensures that all nodes in one layer are connected to nodes in the next layer.

Convolutional neural networks (CNNs) are another type of neural network that served as foundation for the TERL method (Pereira *et al*. 2020). Originally designed for processing images, CNNs deal with two-dimensional objects formed by pixels. In the case of TERL, they are applied to TE classification by pre-processing and transforming TE sequences (one-dimensional objects) into two-dimensional data (like images) and feeding them into the CNN. The method seeks to create a set of attributes that best represents the data for subsequent classification.

CNNs were also used in the DeepTE (Yan *et al*. 2020) and Inpactor2 (Orozco-Arias *et al*. 2022) tools. DeepTE employs vectors formed by k-mer frequencies as attributes. Eight different models were trained and ordered according to a tree structure to enable TEs classification at different levels such as superfamily and order. One of these trained models was designed to detect domains within the sequence to correct misclassification. Inpactor2 uses different CNNs to identify LTR-RT in large genomes and classify them at the superfamily and family levels, showing promising results in terms of reduced runtime and accuracy.

## 3. Interaction Prediction

In conventional supervised learning, the objective is to construct a prediction function based on a set of instances, known as the training set. Each training instance is represented by a vector of attributes and is associated with one or more labels (Pliakos *et al*. 2018). Instances belonging to only one label constitute single-label tasks, while instances belonging simultaneously to two or more labels constitute multi-label tasks. However, this is not the only way to represent data.

Interaction data is an example of complexly structured data, where there are two sets of instances, each described by its own set of attributes. Bioinformatics encompasses several examples of interaction data, such as gene expression analysis and drug-protein interaction (Pliakos *et al*. 2018). The interaction between two sets can be represented by a bipartite graph and an interaction matrix. Each row and column of the matrix is associated with the instances of each set, and the cells are filled with binary values that represent the existence or absence of interactions between the considered instances (Santos *et al*. 2019). Mathematically, let there be two sets of horizontal instances *H* = {**h**_1_, …, **h**_|*H*|_} and vertical instances *V* = {**v**_1_, …, **v**_|*V* |_} of the interaction matrix **Y**, where each instance is represented respectively by attribute vectors **h**_*i*_ and **v**_*j*_. Every label y(i,j) of the matrix **Y** equals 1 if there is an interaction between instances **h**_*i*_ and **v**_*j*_, and zero otherwise. Figure 1 illustrates the representations for such data structure. In the interaction prediction task, given two sets of instances *H* and *V*, each represented by their own set of attributes, the general objective is to predict whether there is an interaction between each pair of instances in the network. Therefore, the task is to define a function *f* : *H × V* → 0, 1 applicable to unknown (test) instances (Pliakos *et al*. 2018).

**Figure 1.**
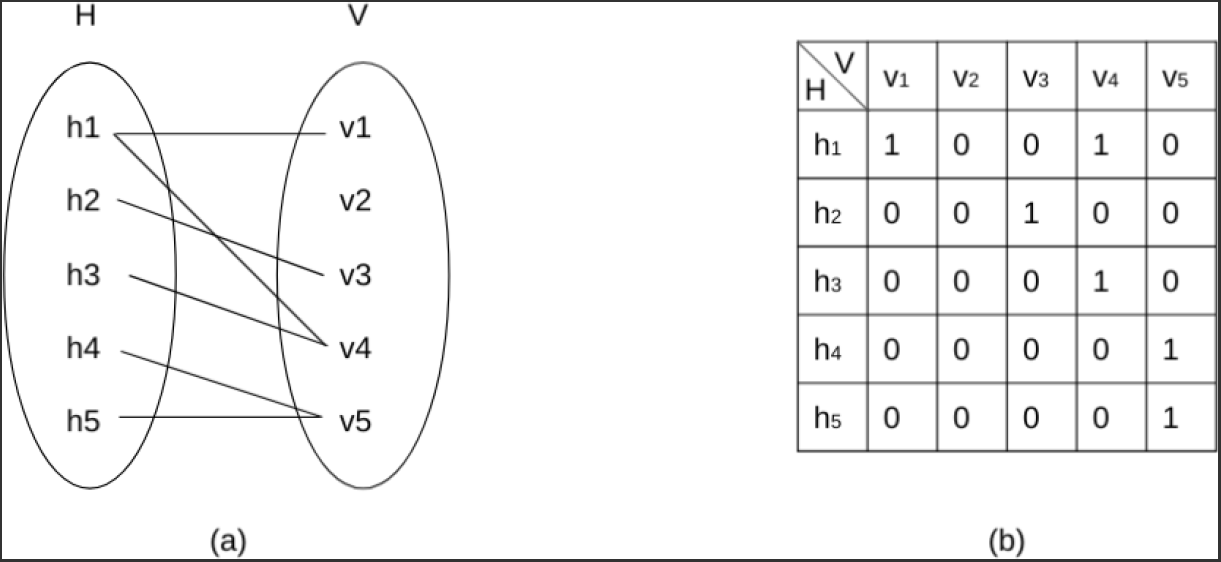
Representation of interaction data. (a) Bipartite graph; (b) interaction matrix. Adapted from Santos *et al*. (2019).

The main challenges of this task, according to Pliakos *et al*. (2018), are computational efficiency and predictive accuracy.

Interaction prediction can be divided into two main categories (Park & Marcotte 2012): prediction applied to a pair of completely unknown instances and prediction applied to a pair where one of the instances is included in the learning process. When interaction prediction is treated as a supervised learning process, it can be divided into three subgroups:

- When interaction prediction is performed on an interaction matrix formed by horizontal instances that were included in the learning process and unknown (test) vertical instances;
- When interaction prediction is performed on an interaction matrix formed by vertical instances that were included in the learning process and unknown (test) horizontal instances;
- When interaction prediction is performed on an interaction matrix formed by unknown (test) horizontal and vertical instances.

Additionally, interaction prediction can be solved using global or local-based methods. The global method adapts traditional algorithms or data so that a single classifier is capable of solving the prediction task. On the other hand, the local method deals with each set of instances (horizontal and vertical) independently, and the results are gathered at the end of the process.

The global single output method concatenates the attribute vectors so that the resulting vector for a pair *k* is in the form **r**_*k*_ = {**h**_**1**_, **h**_**2**_, …, **h**_**n**_, **v**_**1**_, **v**_**2**_, …, **v**_**m**_}. The function *f* is constructed on this attribute space and outputs an interaction vector **Y** of length |*H*| ∗ |*V* |. In the case of decision tree learning, the global method involves constructing a tree from all pairs of training instances. Each split (branch) of the tree occurs based on one of the attributes of a given instance, whether it is horizontal or vertical, and such a split generates the partitioning of the label space (the vector **Y**). The goal of each split is to reduce variance within each partition. Therefore, each leaf of the resulting tree is associated with a subset of the interaction matrix.

In the local method, the interaction prediction of subgroups is explored. Two decision tree models are constructed: the first one is based on known horizontal instances and aims to predict interactions in subgroup 2, while the second is based on known vertical instances and aims to predict interactions in subgroup 1. In the final step of the method, the predictions of the two models are combined, and the final result is presented.

The model used in this paper adapts the global method in the context of decision trees so that data transformations (such as concatenation of attribute vectors) are not required, and the output is multi-label. The so-called global multiple output allows for the direct application of the method to interaction data, and correlations between the sets of instances *H* and *V* are explored. The next section details the global-based method used in our experiments.

## 4. Predictive Bi-Clustering Trees

A Predictive Clustering Tree (PCT) is a decision tree constructed using a top-down approach, in which the root node corresponds to the entire set of instances and each derived node corresponds to a subset or cluster. The clusters are recursively divided into other nodes according to the tests applied to one of their attributes. To determine the test that best divides a cluster, all attributes and their respective values are considered. The goal is to define tests that reduce the variance between the labels of the instances of each generated cluster. The variance of a cluster *S* is defined as:

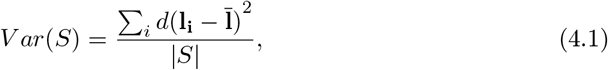

where *d* is the distance between each label vector **l**_**i**_ and the average label vector 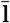 of cluster *S*.

A Predictive Bi-Clustering Tree (PBCT) is a specific case of a PCT that trains a decision tree considering two sets of instances and their respective attributes: the horizontal instances *H* and the vertical instances *V*. As explained in Section 3 for the interaction prediction task, the two sets form a binary interaction matrix as illustrated in Figure 1. The matrix’s rows denote the label vectors of instances within set *H*, while its columns are the label vectors of instances within set *V*. In both scenarios, these label vectors symbolize interactions between row and column instances. A PBCT aims to predict whether there is interaction or not between each pair of instances considered, thus the constructed function is *f* : *H × V* → 0, 1.

The definition of the tests applied to each node in a PBCT is similar to that of a PCT, and the criterion is the same (minimizing the variance within the generated clusters). However, attributes from both dataset *H* and dataset *V* can be used in the tests at the internal nodes of the tree. As a result, labels in Equation 4.1 can correspond to either horizontal or vertical instances. Figure 2(b) illustrates the PBCT method, where *ϕ*_*H,i*_ and *ϕ*_*V,j*_ represent, respectively, the test application referring to an attribute of the horizontal and vertical instances of the interactions matrix. When an attribute from dataset *H* is used in the node’s test, the interaction matrix is partitioned horizontally. Similarly, when an attribute from dataset *V* is used, the matrix is partitioned vertically. Thus, the label space can be partitioned in two directions during the tree learning process, which dynamically considers their correlations. These partitions are represented in Figure 2(c). To determine the partitioning between vertical and horizontal instances, the reduction in variance generated by the split is computed for each row or column of the matrix. In the case of a horizontal split (*ϕ*_*h,i*_), the variance of the resulting subset (*S*) is calculated as

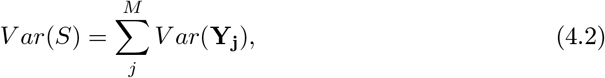

where *M* is the number of columns in the matrix. Similarly, in the case of a vertical split (*ϕ*_*v,j*_), the variance of the resulting subset is calculated as

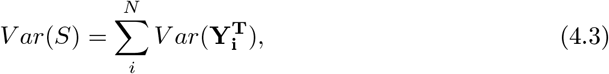

where **Y**^**T**^ is the transpose of the matrix **Y** and *N* is the number of rows. Therefore, the heuristic *h* has the final form of the Equation 4.4, where the terms 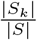 and 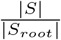 prevent partitions from being split in only one direction. The process of building the tree is presented in Algorithm 1.

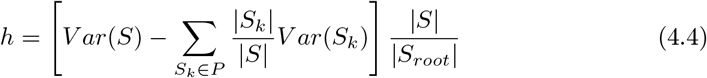

**Figure 2.**
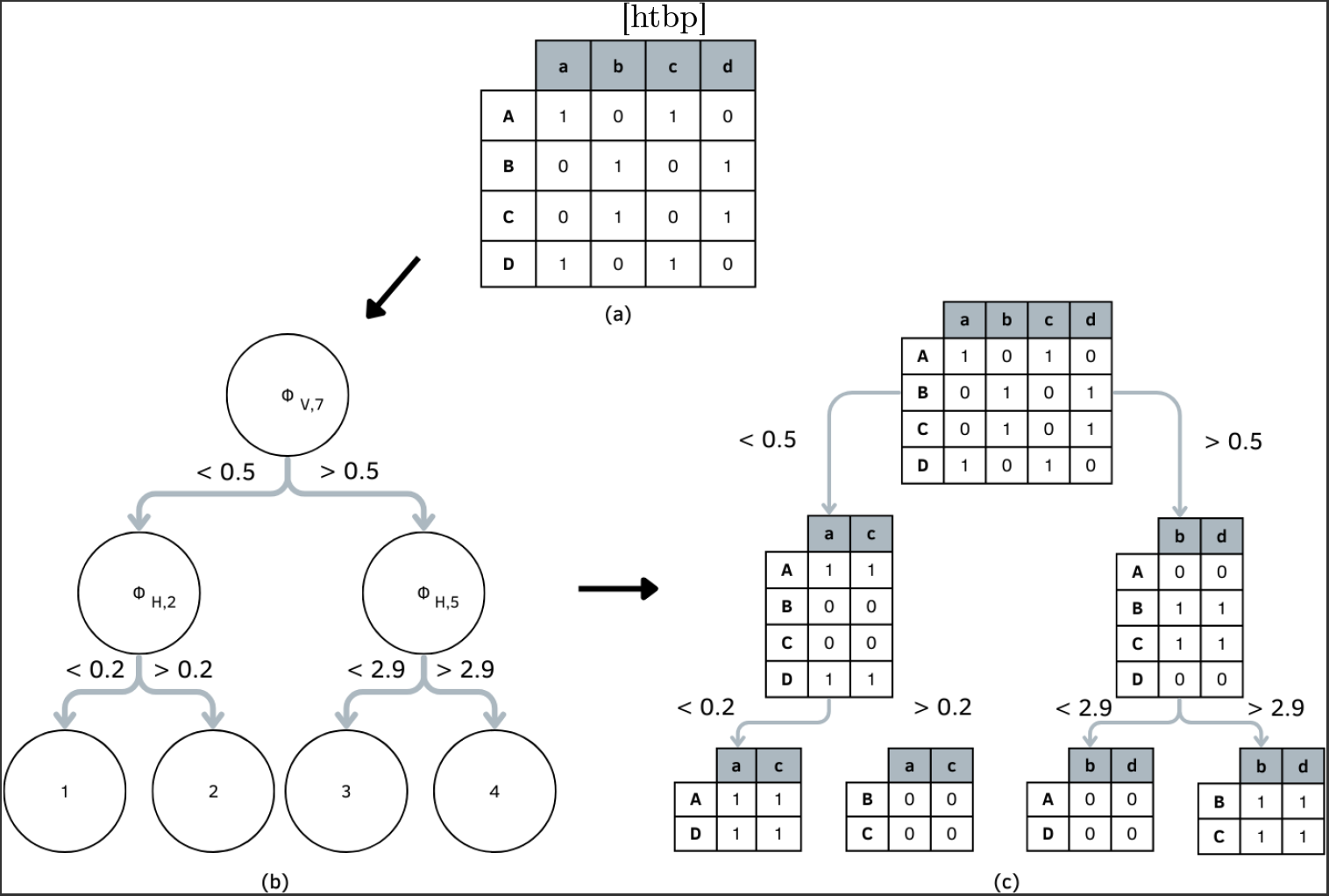
Illustration of the PBCT learning process: (a) interaction matrix; (b) decision tree; (c) corresponding partitioned interaction matrix. Figure adapted from (Pliakos *et al*. 2018).

## 5. LTR Retrotransposons classification as an Interaction Prediction Problem

Our proposal uses the following conventions: the sequences of LTR retrotransposons represent the horizontal instances in the interaction matrix: *H*_*L*_ denotes training (learning) instances and *H*_*T*_ denotes test instances. The conserved protein domains represent the learning vertical instances, denoted as *V*_*L*_.

### Algorithm 1

PBCT learning process. Adapted from (Pliakos *et al*. 2018).

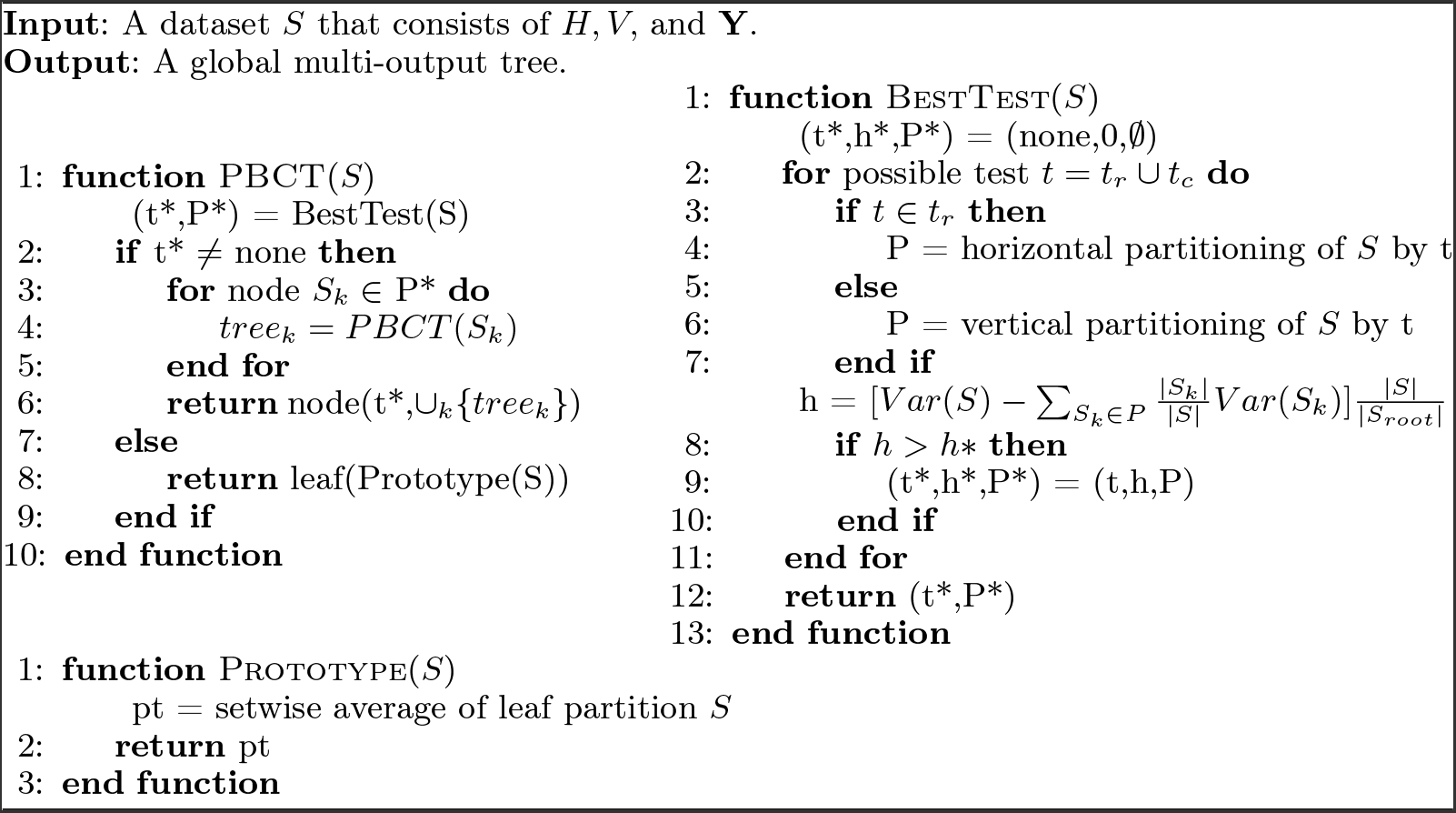

The datasets used in this work allow the construction of a non-diagonal interaction matrix, where the labels correspond to the interactions between LTR retrotransposons and conserved protein domains. These labels can therefore be interpreted as indicating the presence or absence of conserved protein domains within the analyzed TE sequences. This selection is based on the fact that each superfamily of LTR retrotransposons has a distinct set of protein domains, and their identification within a sequence enables accurate classification at the superfamily level. This connection between interaction prediction and classification tasks is significant.

Additionally, the conserved domains data constitute a training set *V*_*L*_ and are exclusively used to build the classifier, while the test set comprises only attributes related to LTR retrotransposon sequences **H**_**T**_. The rationale for this choice is elaborated in Subsection 5.2.

### 5.1. LTR Retrotransposons

The model proposed in this work was evaluated using two organisms: *Drosophila melanogaster* and *Arabidopsis thaliana*. Therefore, datasets were constructed for each organism.

The TE sequences dataset was constructed based on the data obtained by Schietgat *et al*. (2018) and follows similar constructions for both the model’s training and testing sets. In their work, transposable element sequences were extracted from the public database Repbase (Jurka *et al*. 2005). The sequences related to the target organism were excluded, and the considered attributes were the conserved protein domains identified within these sequences. The RPS-Blast (Reverse Position-Specific Blast) tool (Camacho 2017) was employed to identify the domains present in the LTR retrotransposons sequences. Figure 3 provides an illustration of how a TE candidate sequence was represented in (Schietgat *et al*. 2018). For each candidate sequence, the included information indicates which domains were identified in it, the positions at which they were found, and the confidence level of this information (e-value).

**Figure 3.**
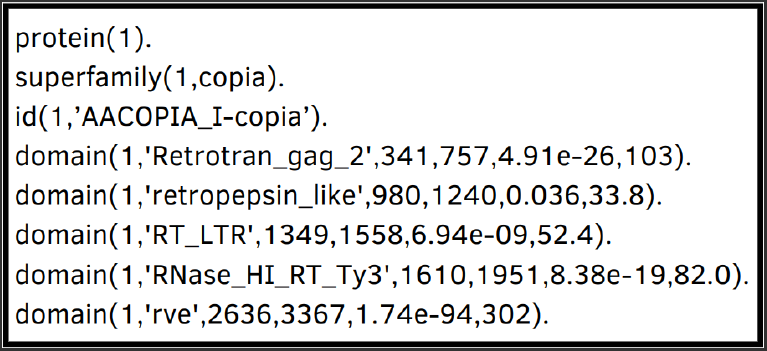
Example of LTR retrotransposon sequence represented by the its identified conserved protein domains. Each domain contains the sequence number, superfamily identification, sequence ID, the conserved domain, its predicted start and end positions, and the associated e-value of the prediction. Extracted from Schietgat *et al*. (2018), Figure 3, p.7.

In contrast to (Schietgat *et al*. 2018), this work considers pseudo components as attributes of the LTR retrotransposon sequences. Pseudo components are attribute vectors resulting from discrete models that analyze the sequences and express their main pattern information. To extract the pseudo components, the dataset created in (Schietgat *et al*. 2018) was used to retrieve the original LTR retrotransposon sequences.

In this work, as in (Schietgat *et al*. 2018), it is considered that each identified LTR retrotransposon sequence starts at the first position of the first identified domain and ends at the last position of the last domain. For example, the sequence in Figure 3 starts at position 341 and ends at position 3367 of the candidate sequence 1. Pse-in-One 2.0 (Liu *et al*. 2017) was used to generate the attribute vectors (pseudo components) of the sequences. Table 1 shows the number of sequences in the training and test datasets for each organism and superfamily.

**Table 1.**
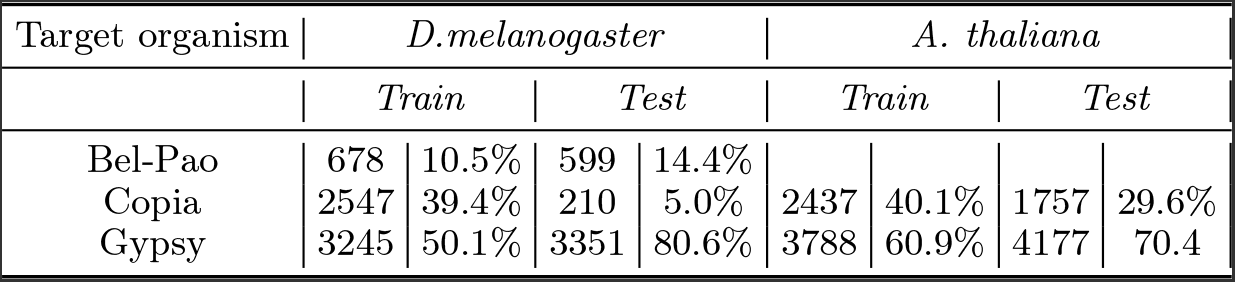
Number of sequences in the training and testing datasets for each organism (*D. melanogaster* and *A. thaliana*) and superfamilies (Copia, Gypsy, and Bel-Pao)

### 5.2. Conserved protein domains

The dataset of protein domains present in the TE sequences was also constructed based on data obtained by Schietgat *et al*. (2018). However, for this dataset, the data in (Schietgat *et al*. 2018) was used to recover the sequences of the domains themselves. As an example, let us return to Figure 3, but now there are five sequences to be recovered, each corresponding to a different domain: Restrotrans_gag starting at position 341 and ending at 757; retropepsin_like starting at position 980 and ending at 1240; and similarly for the other domains.

Therefore, sequences representing protein domains contained in the considered LTR retrotransposons were obtained. From these, consensus sequences were constructed for each protein domain, and the attributes considered are pseudo components of these consensus sequences. As an example, let A and B be two different sequences of Copia superfamily LTR retrotransposons such that sequence A has the Retrotrans_gag domain between positions 50 and 350, and sequence B also has this domain, but between positions 80 and 420. Although it is the same domain, the sequences representing it are different. As the goal is to obtain attributes that represent the Retrotrans_gag domain, a consensus sequence is created from sequences A and B, and the pseudo component vectors are extracted from it.

The creation of the consensus sequence for each domain was performed according to the following process: the MAFFT tool (Katoh *et al*. 2002) was used to perform the multiple alignment of sequences; from this alignment, the HMMER tool (Johnson *et al*. 2010) was used to create the HMM profile; and from this profile, using HMMER, the consensus sequence was created. The objective with this is to have an attribute vector for each conserved domain.

The resulting interaction matrix is illustrated in Figure 4, where the rows are associated with LTR retrotransposon sequences belonging to one of the three superfamilies, and the columns are associated to consensus sequences of domains for each superfamily. As introduced in Section 3, the rows constitute the set of horizontal instances H and the columns the set of vertical instances V. The matrix is filled with binary values, where the value 1 means that the corresponding domain is present in the considered LTR retrotransposon sequence, and 0 means it is not.

**Figure 4.**
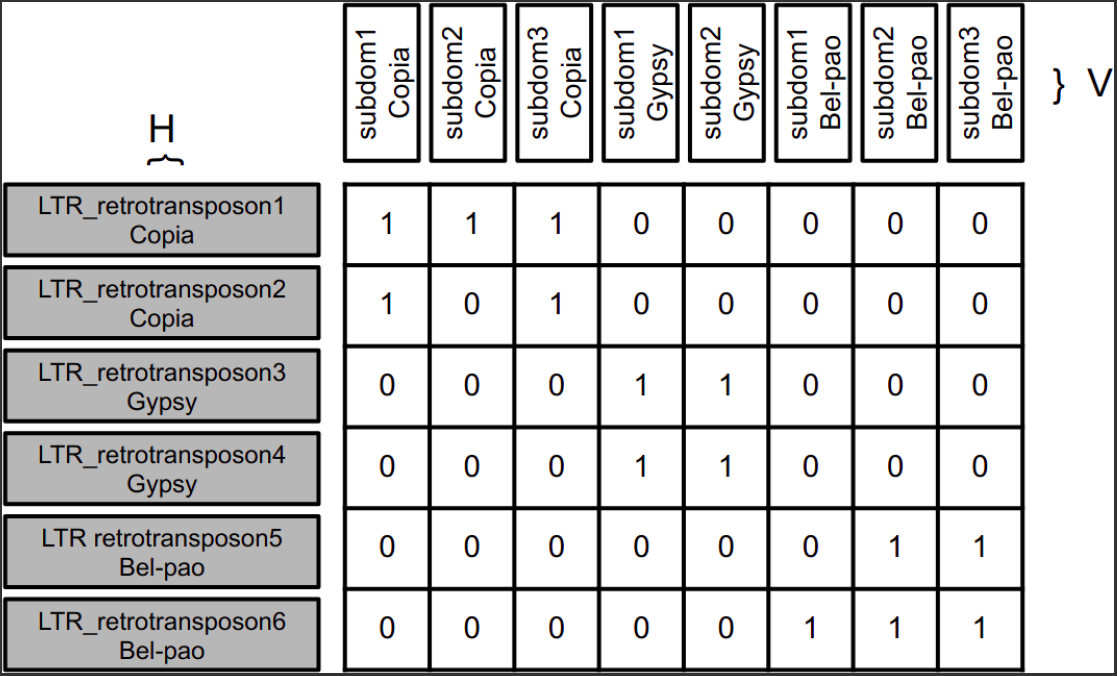
Interaction matrix in which rows are associated with LTR retrotransposon sequences, and columns with the consensus sequences of the protein domains present in each superfamily.

This approach for constructing *V*_*L*_ attributes cannot be applied in the testing scenario. This limitation arises because the feature extraction process relies on the presence of a consensus sequence for each domain present in the training dataset. For a test sequence we do not have these consensus sequences. Thus, for a test TE candidate sequence, the PBCT is used to predict the interactions between the TE sequence and the conserved domains (represented by its consensus sequence) obtained in the training process.

## 6. PBCTs for Classifying LTR Retrotransposons

Following the dataset creation process outlined in Section 5, the PBCT method can be applied to predict interactions between LTR retrotransposons and conserved protein domains. These predictions can subsequently be used for classifying the LTR sequences at the superfamily level.

Figure 5 illustrates the process of interaction prediction for an unknown LTR retrotransposon sequence from *H*_*T*_ (x) with protein domains (sDP). While traversing the tree, when the test relates to an attribute of an LTR retrotransposon (horizontal) (*ϕ*_*H,i*_), the pair proceeds along a specific branch of the tree based on the test result. However, when the test pertains to an attribute of a protein domain (*ϕ*_*V,j*_), both branches of the tree are followed. This happens because when testing a candidate TE we do not have access to its protein domains. Thus, we use the tree to predict the protein domains which interact with it. As a result, instances may be classified into more than one leaf of the tree.

**Figure 5.**
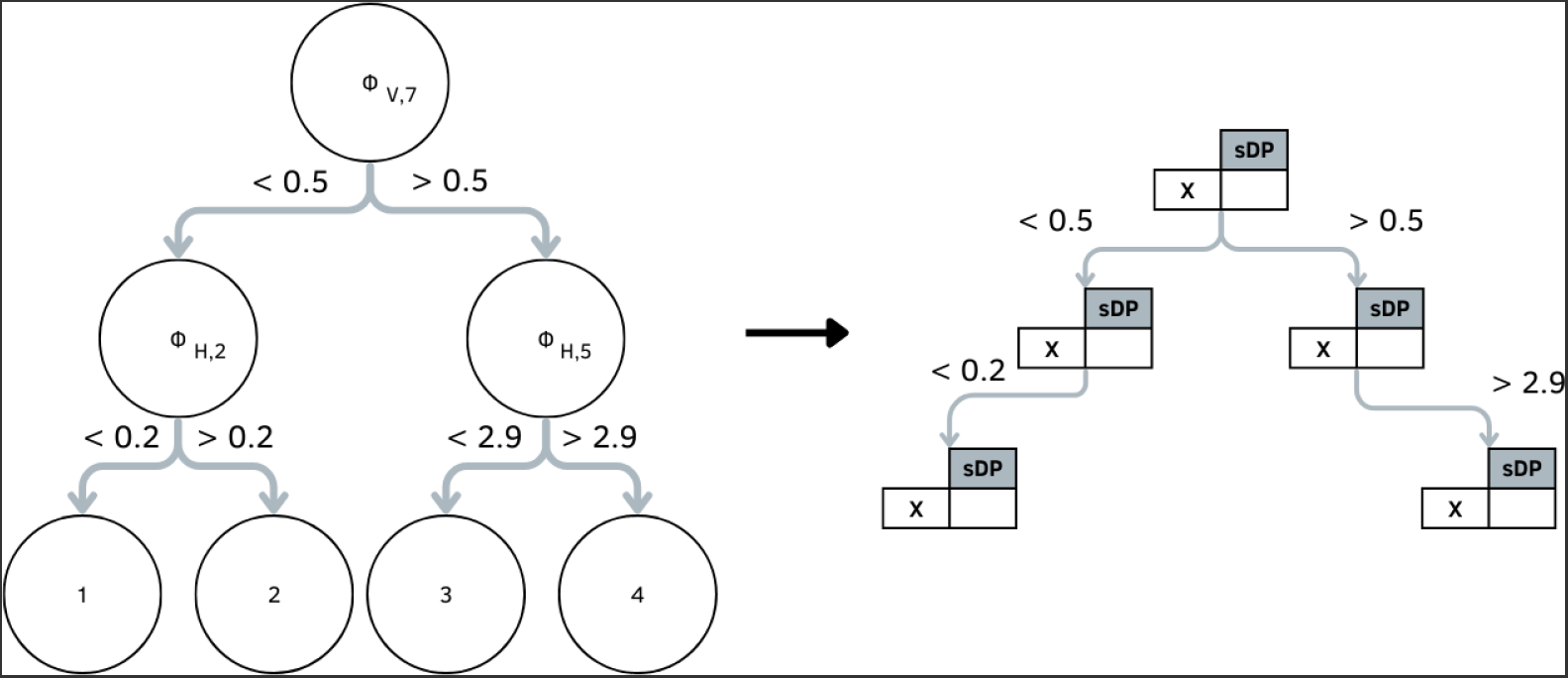
Illustration of the PBCT prediction process through the proposed method.

Each leaf contains the probability of an interaction existing between the unknown LTR retrotransposon sequence and the conserved protein domain. The final prediction for each instance involves retrieving all leaf nodes associated with the TE, along with their prediction values, and then calculating their average. This process is illustrated in Algorithm 2. Since the domains that can be found in each superfamily are different, it is possible to calculate the probability of the sequence belonging to each superfamily based on the probability vector provided by PBCT.

### Algorithm 2

Predictions with a PBCT

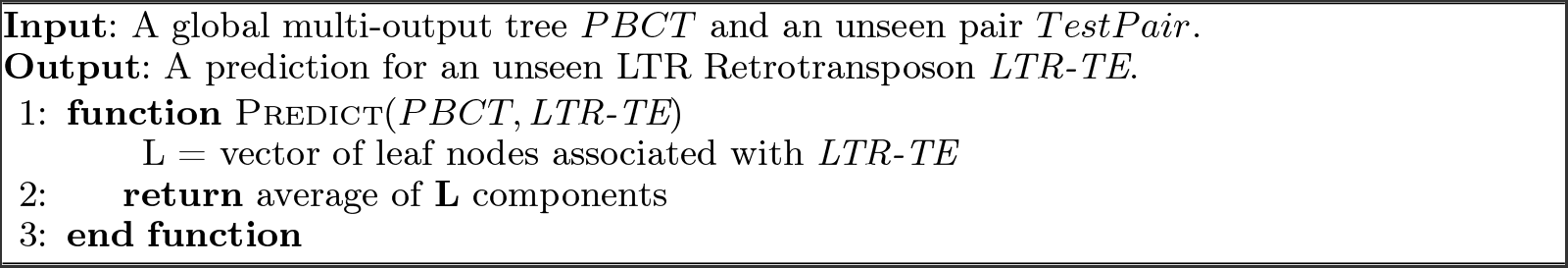

## 7. Experiments and Discussion

### 7.1. Evaluation Measures

Our proposal was evaluated using precision-recall (PR) curves for each superfamily. The choice of these curves is justified by the fact that the primary objective of the classifier is to identify true positive cases, i.e., the candidate sequences that genuinely belong to a superfamily. Precision represents the percentage of correct predictions, while recall represents the percentage of annotations that were correctly predicted. The PR curves were calculated individually. Therefore, true positive (TP) cases occurred when the model correctly predicted the superfamily, false positives occurred when the predicted superfamily was incorrect, and false negatives occurred when the model incorrectly predicted a different superfamily than the actual one. A PR curve plots the precision of the model as a function of its recall Schietgat *et al*. (2018). Equations 7.1 and 7.2 present the formulas for Precision (P) and Recall (R), respectively.

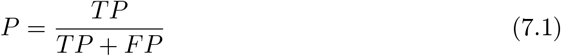

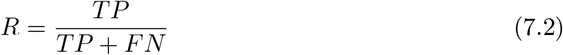

As previously mentioned, the trained model predicts the probability of each sequence having or not the protein domains considered for each superfamily. The classification at the superfamily level is performed in a subsequent analysis, where the superfamily with the highest number of identified domains is predicted.

During the model evaluation, a threshold value *t* is used to determine the superfamily in which a sequence is classified. Each specific threshold value corresponds to a single point on a PR curve obtained by varying *t* from 1.0 to 0.0. Decreasing the threshold results in an increase in the number of sequences classified as positive, thereby increasing recall while precision may either decrease or increase (tending to decrease). In this analysis, each sequence is predicted among the superfamilies Copia, Gypsy, Bel-Pao (only for the organism *Drosophila melanogaster*), or None. A sequence is classified as None when there is a tie in the number of predicted domains for each superfamily, causing the classification to be indecisive between two or three superfamilies. The sequences predicted as None are not considered for the prediction calculation but are included in the recall calculation.

Another measure used for evaluating the created models was the F1 score, as presented in Equation 7.3. F1 is a metric that computes the harmonic mean of precision and recall, indicating the efficiency of classifying TEs when precision and recall are equally relevant.

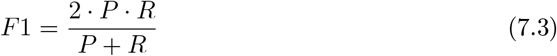

In (Schietgat *et al*. 2018), the obtained results are compared to the three most commonly used methods in the tasks of TE identification or classification at the superfamily level: RepeatMasker, Censor, and LtrDigest. We also compare our proposal with the results of these three methods. We use the same results provided by Schietgat *et al*. (2018). Additionally, a baseline is also used in the comparisons, which makes predictions based only in the presence of one key protein domain predicted by RPS-Blast (Schietgat *et al*. 2018).

### 7.2. Results and Discussion

In total, 4160 sequences from the organism *Drosophila melanogaster* and 5934 sequences from the organism *Arabidopsis thaliana* were classified by the model during the tests. The creation of all decision trees followed the default PBCT hyperparameters, which are as follows: undefined maximum tree depth, minimum number of instances in leaves set to 20, minimum quality of each node set to 0, and maximum leaf variance set to 0.

Various pseudo components were considered for attribute extraction from the sequences. We present here the results for the attributes that exhibited the best overall performance. The criterion used to evaluate these performances was the F1-score of each model. Additionally, the adoption of the F1-score aligns with the approach used in (Schietgat *et al*. 2018), ensuring consistency in the assessments. Based on the established criterion, the most effective pseudo components are DACC (Liu *et al*. 2017) for the organism D. melanogaster and Mismatch (Liu *et al*. 2017) for the organism

A. thaliana. DAAC is an autocorrelation feature, standing for dinucleotide-based auto-cross covariance and based on the 148 physicochemical indices of dinucleotides Dong *et al*. (2009). Mismatch is a features containing nucleic acid composition. It counts the occurrences of kmers, allowing at most m mismatches Leslie *et al*. (2004).

#### 7.2.1. PBCT predictions for each Superfamily

Since PBCT output predictions for all superfamilies, we initiate our analysis comparing the precision-recall curves generated for each superfamily individually. Figure 6 illustrates these curves for the organism *D. melanogaster*. By their overall shape, it is evident that PBCT faced challenges in classifying the Bel-Pao and Copia superfamilies. For the latter, recall reached a maximum value of 0.72, but precision did not exceed 0.2 at any threshold. In the case of Bel-Pao, the peak precision at 0.84 rapidly declines, preventing recall from reaching the 0.2 mark. Figure 7 shows the curves for the organism *A. thaliana* regarding predictions for Copia and Gypsy superfamilies (*A. thaliana* does not contain Bel-Pao). In this case, the Copia curve exhibits greater stability and reaches a maximum precision value of 0.31 and a recall value of 0.83.

**Figure 6.**
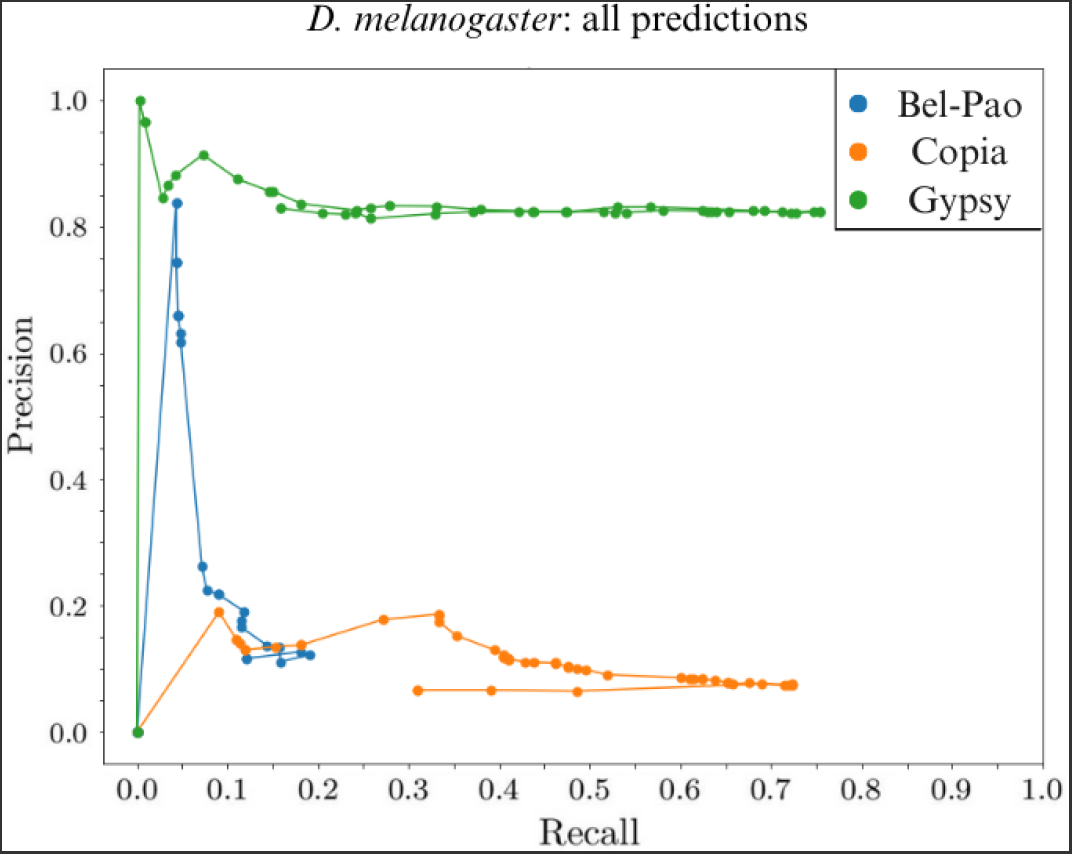
PBCT precision-recall curves for all superfamilies (*D. melanogaster*).

**Figure 7.**
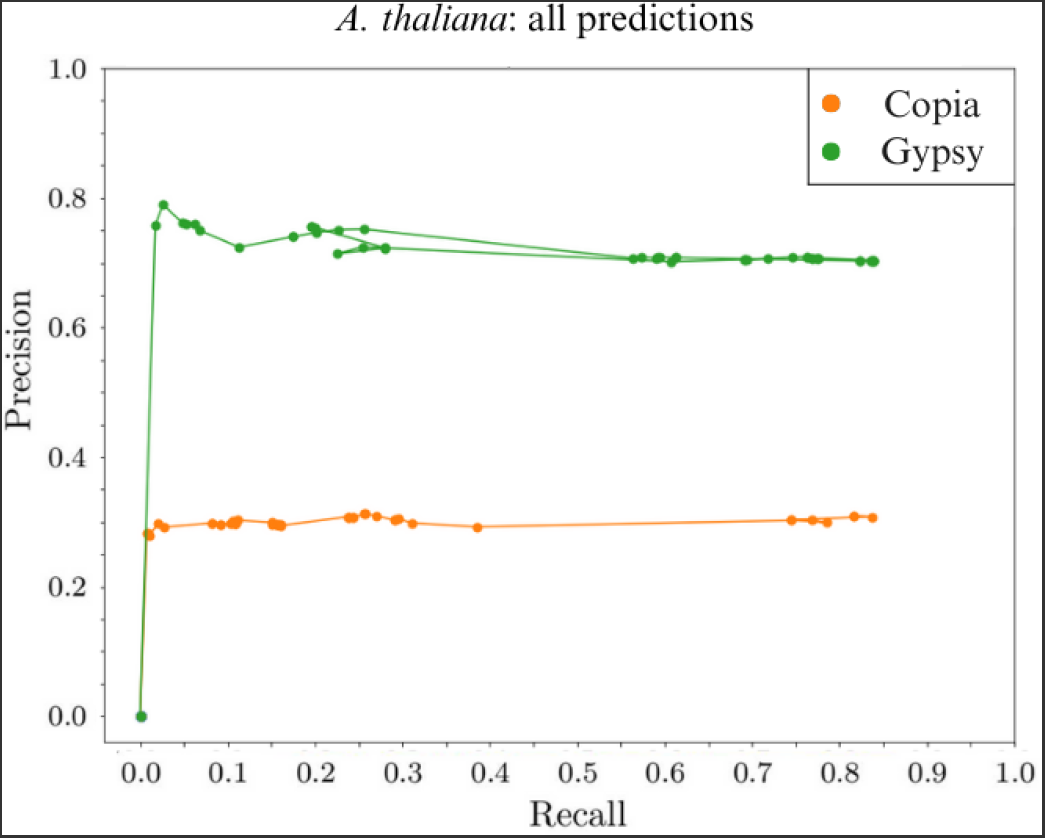
PBCT precision-recall curves for all superfamilies (*A. thaliana*).

For both *A. thaliana* and *D. melanogaster*, the Gypsy curves reach the highest values for precision and recall. In the case of *D. melanogaster*, the maximum recall of 0.82 is accompanied by a precision of 0.75. In the case of *A. thaliana*, the maximum recall of 0.84 is achieved with a precision of 0.70. Furthermore, it can be observed that each curve exhibits an initial threshold interval in which, as the threshold increases, the recall also increases until reaching a maximum value. After this interval, which varies in length depending on the superfamily, recall follows the typical behavior: it decreases as the threshold increases.

To justify this behavior, it is necessary to remember that each sequence is annotated with the superfamily that has the highest fraction of predicted domains. From this, it can be said that this atypical behavior of the curve is due to the way superfamily predictions are made. Since PBCT returns the probability of a sequence containing or not a domain, for small thresholds, a significant number of domains are predicted. This increases the chances of ties occurring in the number of predicted domains among different superfamilies. As mentioned in Subsection 7.1, when the fraction of predicted domains is equal between two or more superfamilies, the model becomes indecisive and cannot classify the analyzed sequence, associating it with the “None” class. According to Equation 7.2 and the definition of recall, these sequences constitute false negative cases, which contribute to decrease the recall value.

Another important point is that the interval of increasing recall varies among the curves because the distribution of test sequences in the dataset across superfamilies is not uniform. This could be observed in Table 1. This atypical behavior is attenuated in the case of the organism *A. thaliana* because occurrences of sequences not classified according to the threshold are less frequent and occur in smaller numbers.

Next, the performance of PBCT in each superfamily will be compared to the performances obtained by other classification methods from the literature.

#### 7.2.2. Comparing PBCT with other Classification Methods

Figure 8 presents the precision-recall curves for the Copia superfamily in the organism *Drosophila melanogaster*. PBCT achieves a maximum recall value of 0.72 when it predicts 152 out of the 210 Copia sequences in the dataset. The RepeatMasker, LtrDigest, and baseline models generate predictions with 100% confidence, represented by a single point on the graph. The PBCT curve surpasses the baseline in recall but not in precision. Additionally, the PBCT curve remains below the TE-Learner^*LT R*^ curve at all points on the graph. The points for LTRDigest and RepeatMasker exhibit high values of precision and recall, respectively. The Censor curve mostly lies below the PBCT curve, but shows higher values in precision (0.43 at 0.04 recall) and recall (0.86 at 0.01 precision).

**Figure 8.**
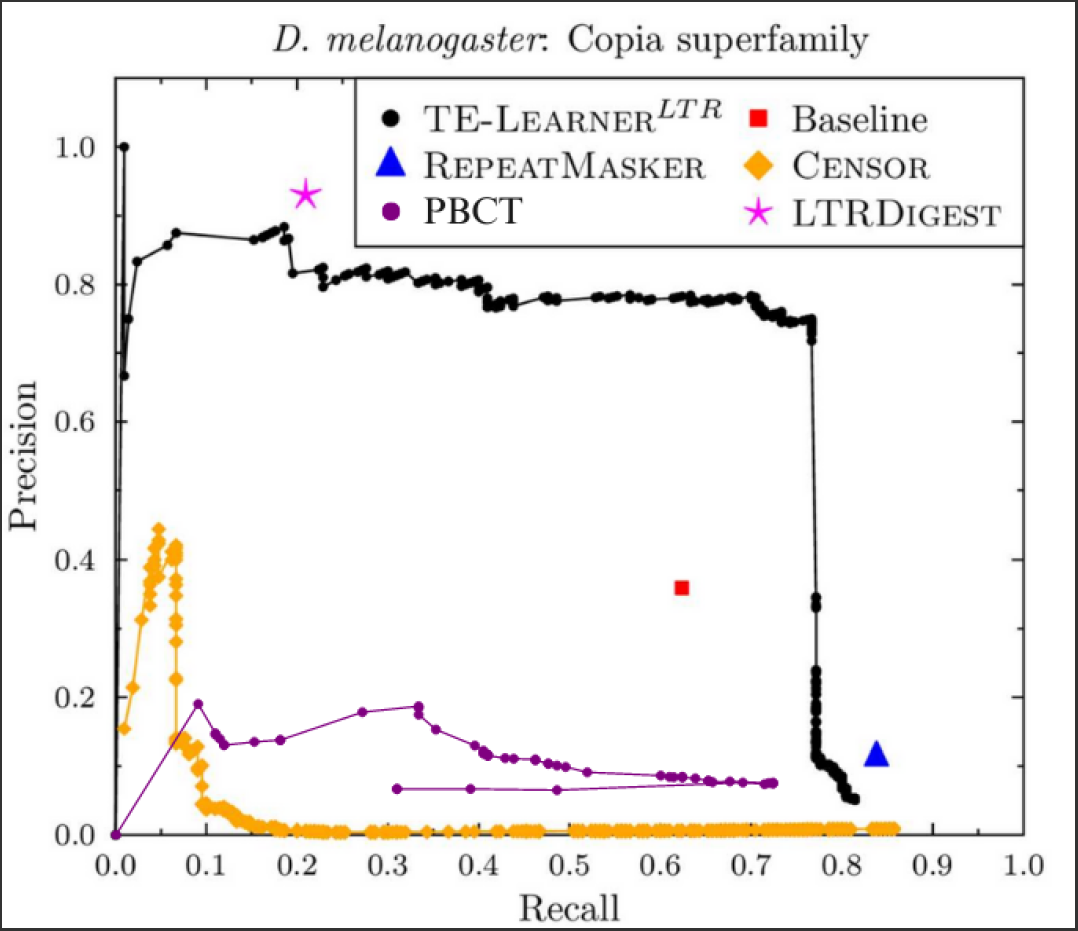
Precision-recall curves for the Copia superfamily (*D. melanogaster*).

Figure 9 displays the curve for the Copia superfamily in the organism *A. thaliana*. Both recall and precision values are higher than those for *D. melanogaster*. Additionally, the maximum recall value achieved (0.83) is higher than any other method, especially

**Figure 9.**
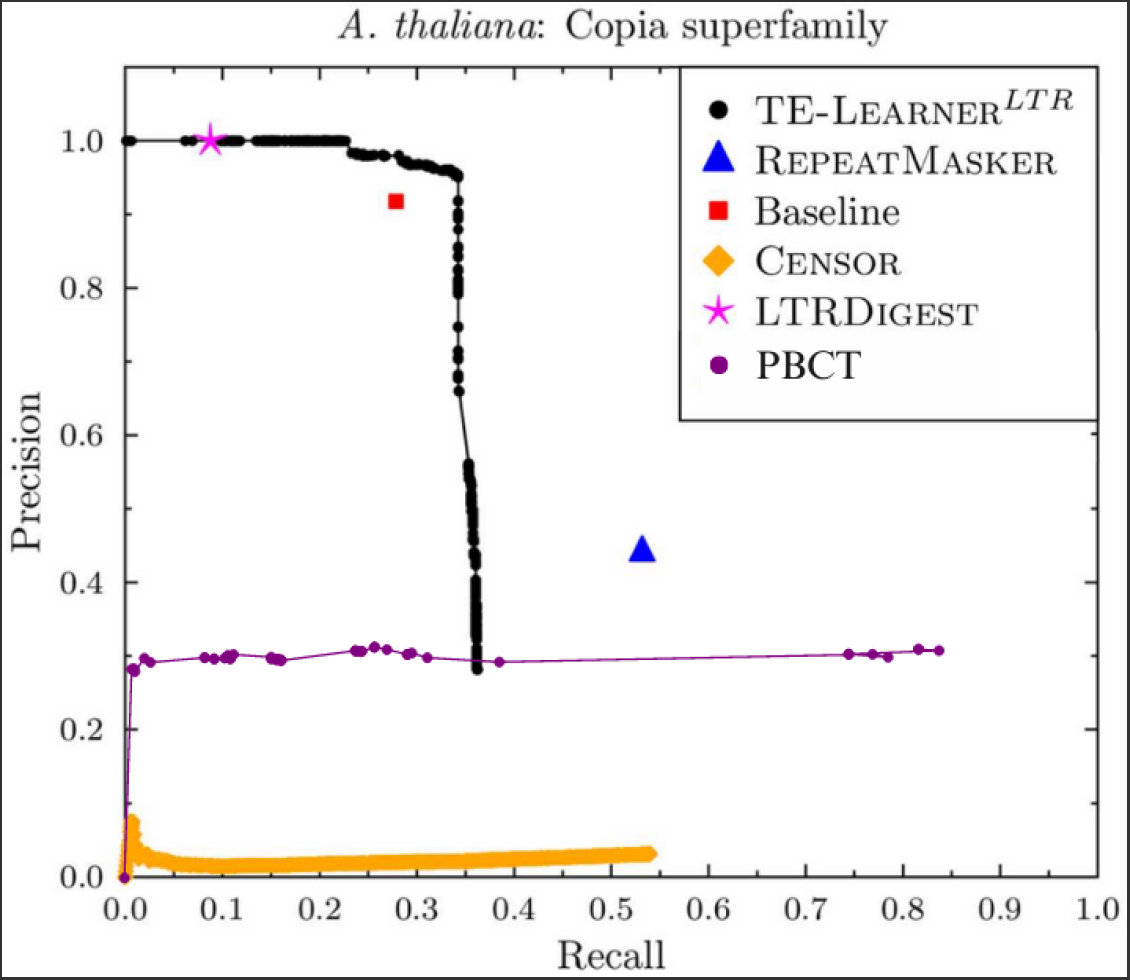
Precision-recall curves for the Copia superfamily (*A. thaliana*).

TE-Learner^*LT R*^, which has a maximum recall near 0.35 at the expense of precision. For PBCT, although its maximum precision of 0.31 is only higher than the Censor curve (which exhibits very low values), the increase in recall is not accompanied by a decrease in precision.

Figure 10 presents the results for the Gypsy superfamily in the organism *D. melanogaster*. Here, the PBCT curve remains above the other methods in almost all points, except for one point where the TE-Learner^*LT R*^ curve has a slight advantage in precision. PBCT achieves a maximum precision of 1.0 at a recall of 0.003. Both RepeatMasker and Censor achieves their highest recall values of 0.80. However, unlike them, PBCT can increase recall without sacrificing precision, as its maximum recall of 0.75 occurs at a precision of 0.82.

**Figure 10.**
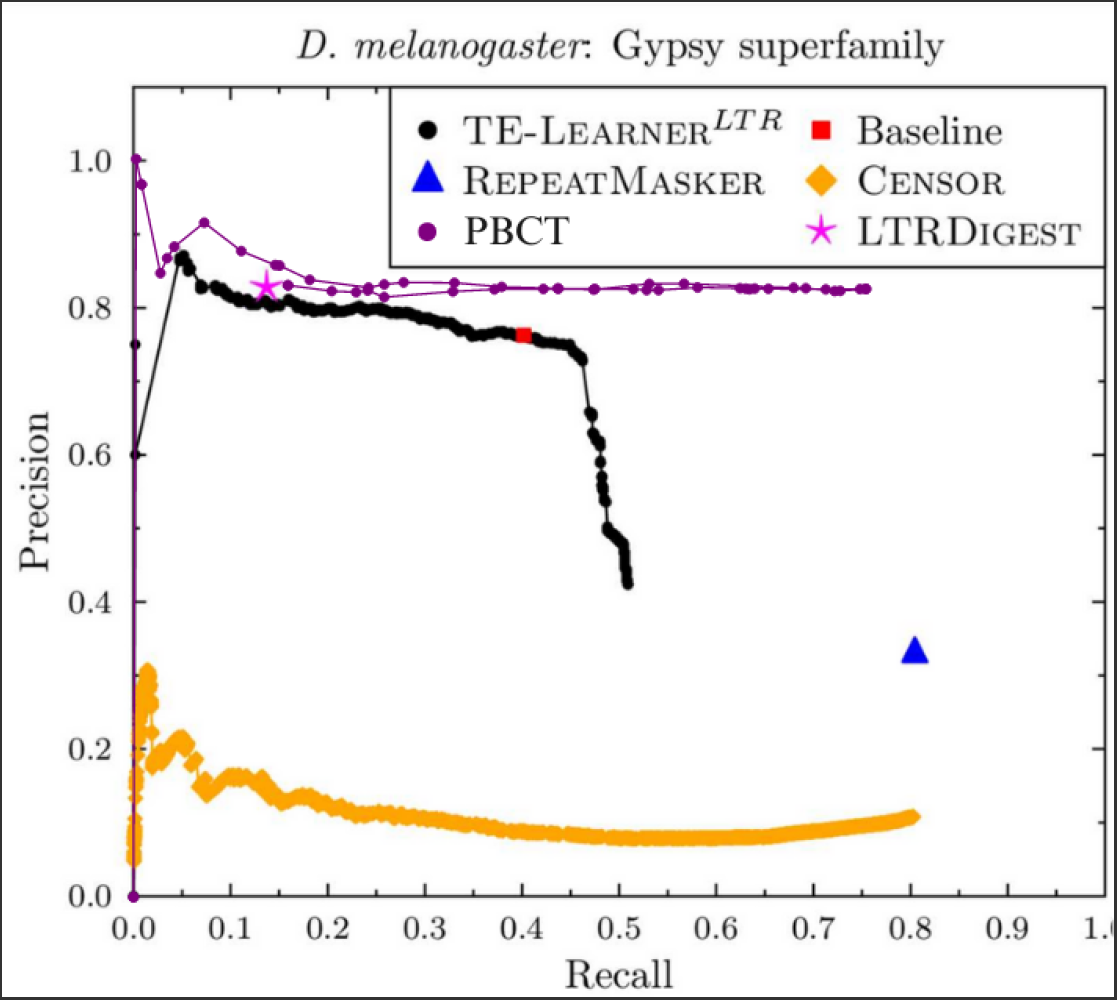
Precision-recall curves for the Gypsy superfamily (*D. melanogaster*).

The precision-recall curves for the Gypsy superfamily in the organism *A. thaliana* are depicted in Figure 11. Despite TE-Learner^*LT R*^, LTRDigest and the baseline exhibiting high precision values, once again PBCT is the only method whose curve is able to keep precision while increasing recall. PBCT achieves a maximum recall of 0.82 with a precision of 0.72.

**Figure 11.**
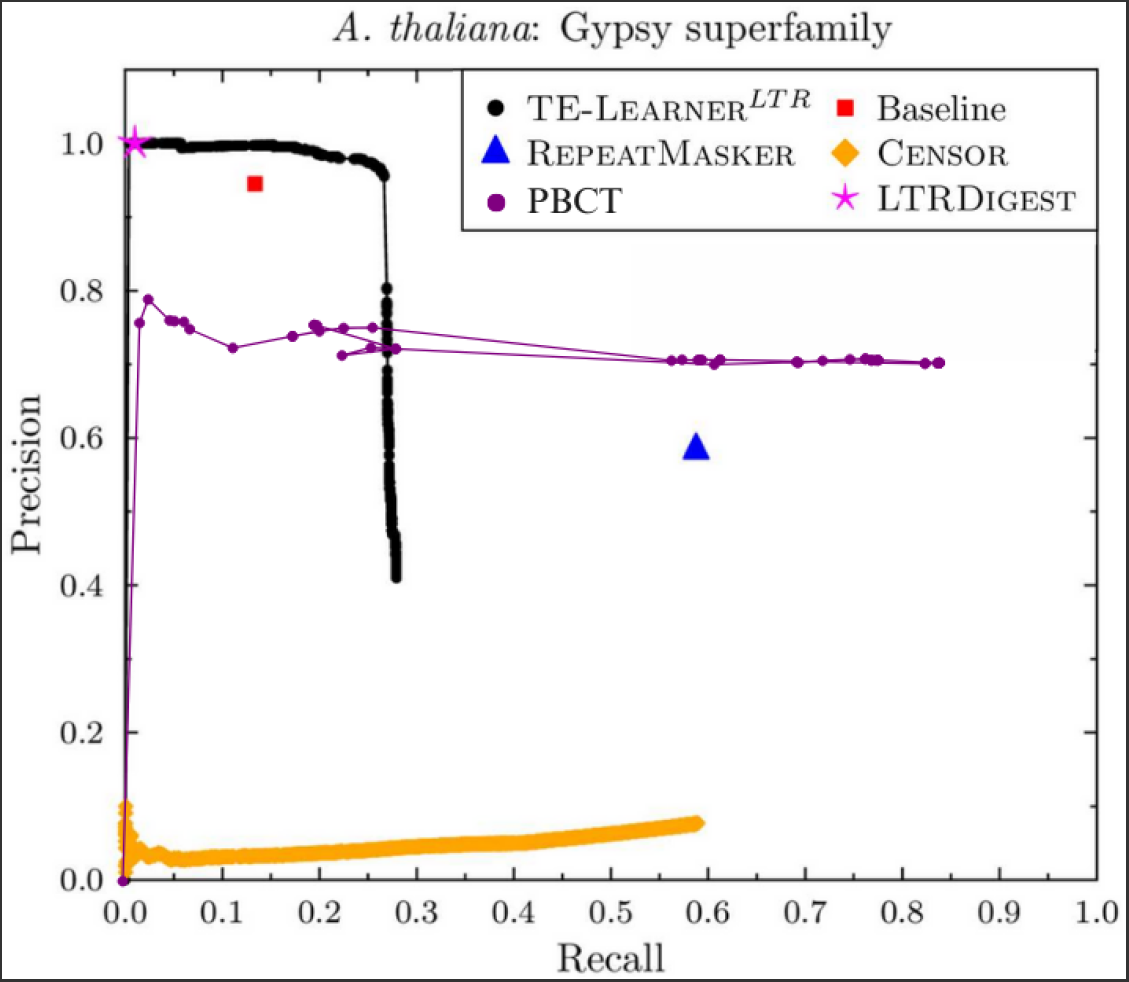
Precision-recall curves for the Gypsy superfamily (*A. thaliana*).

The results for the Bel-Pao superfamily are shown in Figure 12. Remember that Bel-Pao is only present in the organism *D. melanogaster*. The PBCT curve is below the TE-Learner^*LT R*^ curve, reaching a maximum precision of 0.84 at a recall of 0.19. This precision value is lower than that achieved by LTRDigest, which also has a higher recall. The maximum recall achieved by PBCT is 0.19 at a precision of 0.12.

**Figure 12.**
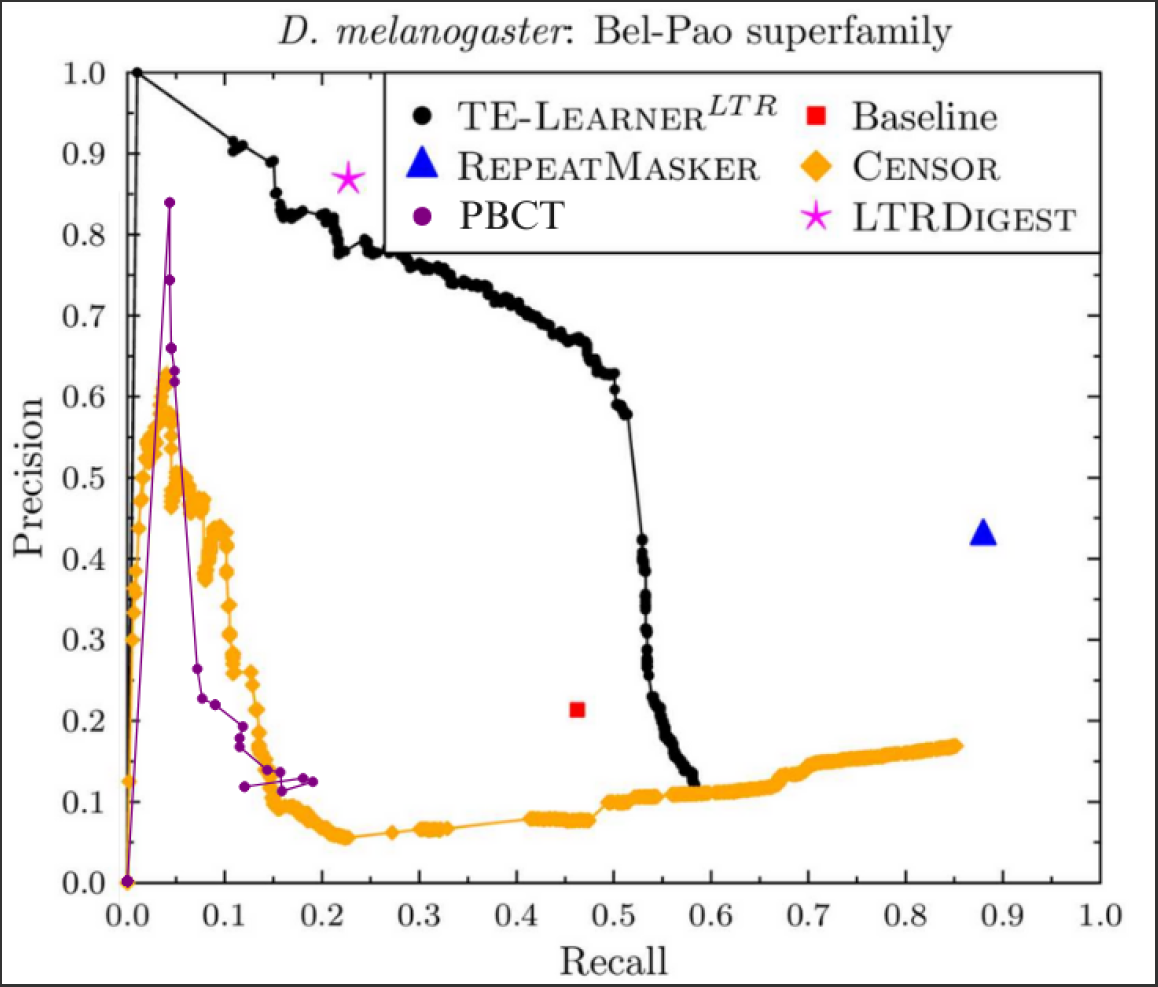
Precision-recall curves for the Bel-Pao superfamily (D.melanogaster).

#### 7.2.3. Discussion

In summary, when compared to TE-Learner^*LT R*^, PBCT shows higher recall values in three out of five classifications. This can be explained by the fact that TE-Learner^*LT R*^ is a framework for sequence identification and classification, so it only classifies the sequences it can identify, limiting its recall. Regarding precision, although the TE-Learner^*LT R*^ achieves higher values, PBCT is more efficient in maintaining a constant precision while increasing recall.

PBCT achieves higher recall values than LTRDigest in all superfamilies (except Bel-Pao). PBCT also outperforms RepeatMasker and Censor in recall in two out of five classifications. As pointed out in (Schietgat *et al*. 2018), both RepeatMasker and Censor are TE identification and classification models that predict a large number of sequences compared to other methods analyzed. As a result, it is expected that their recall reaches higher marks, which, in turn, leads to a decrease in precision. This effect is more clearly observed in the Censor curves, which show rapidly declining precision in four out the five plots.

In general, the performance of PBCT in classifying different superfamilies indicates that the training dataset used caused biases in the classification. This hypothesis is supported by the data in Table 1, which shows that the *D. melanogaster* training dataset contains a proportion of 10.5% Bel-Pao sequences, 39.4% Copia sequences, and 50.1% Gypsy sequences, while the test dataset has sequences in proportions of 14.4%, 5%, and 80.6%, respectively.

This suggests that the model had a greater tendency to classify sequences as Copia rather than as Bel-Pao, as evident in the curves of their respective superfamilies. However, a simple inclination towards classifying most sequences as Gypsy cannot solely explain its PR curve. Notably, it is not just the recall that exhibits high values; precision is also high and tracks the growth of recall. This behavior is consistently observed in all Gypsy curves across all pseudo components used.

Additionally, during tests conducted with balanced datasets, where an equal number of training sequences was used for each superfamily, it was observed that there was no significant improvement in the classification of Copia and Bel-Pao. In contrast, the classification of Gypsy retained the characteristics of the curves presented here, demonstrating high recall accompanied by high precision. This suggests that the conserved domains employed to characterize Gypsy sequences hold substantial significance in distinguishing them from other superfamilies.

In conclusion, Table 2 presents the F1-scores for the analyzed models. Since the results of PBCT, TE-Learner^*LT R*^ and Censor were presented using PR curves, the points of highest F1 along each curve were considered for the analysis.

**Table 2.** F1-score of each method for each target organism and superfamily.

| Target organism | *D. melanogaster* |  |  | *A. thaliana* |  | Average |
| --- | --- | --- | --- | --- | --- | --- |
| Superfamily | Copia | Gypsy | Bel-Pao | Copia | Gypsy |  |
| PBCT | 0.24 | 0.78 | 0.15 | 0.45 | 0.76 | 0.48 |
| TE-Learner<sup>LTR</sup> | 0.76 | 0.57 | 0.56 | 0.50 | 0.42 | 0.56 |
| REPEATMASKER | 0.20 | 0.47 | 0.58 | 0.48 | 0.59 | 0.46 |
| CENSOR | 0.12 | 0.19 | 0.28 | 0.06 | 0.14 | 0.16 |
| LTRDIGEST | 0.34 | 0.24 | 0.36 | 0.16 | 0.02 | 0.22 |

The results show that PBCT performs similarly to LTRDigest, Censor (except for *A. thaliana*), and RepeatMasker in the classification of Copia. For Gypsy, its results are clearly superior to the other methods for both organisms. Concerning Bel-Pao, PBCT performance is the least competitive among all superfamilies. However, on average, PBCT has the second-best performance, being surpassed only by TE-Learner^*LT R*^.

## 8. Conclusions

In this work, we propose to treat the problem of classifying LTR Retrontransposon at the superfamily level as an interaction prediction problem, using Predictive Bi-clustering Trees (PBCTs). The datasets used to train PBCTs included sequence attributes and attributes of the conserved domains characterizing the considered superfamilies.

Our proposal was evaluated using two organisms: *Drosophila melanogaster* and *Arabidopsis thaliana*, with LTR retrotransposon sequences classified into the superfamilies Bel-Pao, Copia, and Gypsy. Our method achieved comparable performances to widely used methods from the literature, particularly through recall and F1 analysis. Furthermore, it obtained the best results in classifying the Gypsy superfamily.

The achieved performance suggests that the application of PBCT to the TE classification task is both feasible and promising. The method can be extended to classify other types of transposable elements and can be applied at different classification levels. Furthermore, there are various aspects of this application that can be explored and studied. For instance, alternative pseudo components can be considered for attribute extraction. Pse-in-One itself offers other models that can be explored, as well as possible combinations between them. To enhance the quality of the created consensus sequences, a filter could be developed to align only sequences with similar e-values. These are all potential research directions that could be pursued to enhance the performance of the model.

## Notes

### Competing Interest Statement

The authors have declared no competing interest.

